# Membrane lipid adaptation of soil Gram-negative bacteria isolates to temperature and pH

**DOI:** 10.1101/2022.10.10.511520

**Authors:** Eve Hellequin, Sylvie Collin, Marina Seder-Colomina, Pierre Véquaud, Christelle Anquetil, Adrienne Kish, Arnaud Huguet

**Affiliations:** Sorbonne Université, CNRS, EPHE, PSL, UMR METIS, F-75005 Paris, France; Muséum National d’Histoire naturelle, CNRS, Unité Molécules de Communication et Adaptation des Microorganismes UMR7245 MCAM, F-75005 Paris, France

**Keywords:** 3-Hydroxy fatty acids, Gram-negative bacteria, soil, temperature, pH, adaptation, proxy, *Bacteroidetes*

## Abstract

3-hydroxy fatty acids (3-OH FAs) are characteristic components of the Gram-negative bacterial membrane, recently proposed as promising temperature and pH (paleo) proxies in soil. Nevertheless, to date, the relationships between the 3-OH FA distribution and temperature/pH are only based on empirical studies, with no work at the microbial level. This work investigated the influence of growth temperature and pH on the lipid profile in three strains of soil Gram-negative bacteria belonging to the *Bacteroidetes* phylum. Even though the non-hydroxy FAs were more abundant than the 3-OH FAs in the investigated strains, we showed the important role of the 3-OH FAs in the membrane adaptation of Gram-negative bacteria to temperature. The strains shared a common adaptation mechanism to temperature, with a significant increase in the ratio of *anteiso* vs. *iso* or *normal* 3-OH FAs at lower temperature. In contrast with temperature, no common adaptation mechanism to pH was noticed, the variations in the FA lipid profiles differing from one strain to another. The models envisioning the reconstruction of environmental changes in soils should include the whole suite of 3-OH FAs present in the membrane of Gram-negative bacteria, as all of them can be influenced by temperature or pH at the microbial level.

## 1 Introduction

Microorganisms (bacteria, archaea and some eukaryotes) are able to adjust their membrane composition in response to the prevailing environmental conditions in order to maintain an appropriate fluidity and ensure the optimal state of the cellular membrane (Hazel and Williams, 1990; Denich et al., 2003; Mykytczuk et al., 2007). Such a strategy is termed ‘homeoviscous adaptation’ (Sinensky, 1974). Bacteria can thus change the number of unsaturations, ramifications or chain length of their membrane lipids – mainly fatty acids – in response to varying environmental conditions (Reizer et al., 1985; Prado et al., 1988; Suutari and Laakso, 1992).

In the same way, the structure of glycerol dialkyl glycerol tetraethers (GDGTs), which are membrane lipids biosynthesized by archaea and some bacteria, is known to be related to environmental parameters (Schouten et al., 2013). In aquatic environments, the relative abundance of the different GDGTs produced by members of the archaeal phylum *Thaumarchaeota* was correlated with the water surface temperature, leading to the development of a temperature proxy (Schouten et al., 2002), the TEX86 index, largely applied to marine and lacustrine paleorecords (Castañeda et al., 2010; Berke et al., 2012). Another type of GDGTs with branched alkyl chains – so-called branched GDGTs (brGDGTs) – were suggested to be produced by bacteria and are ubiquitous in aquatic and terrestrial environments (Schouten et al., 2013). Their analysis in soils/peats (Véquaud et al., 2022) and lake sediments (Martínez Sosa et al., 2021) distributed worldwide showed that their structure varies mainly with air temperature and soil pH, making them increasingly used as temperature and pH paleoproxies. BrGDGTs are the only microbial organic proxies which can be used for temperature reconstructions in both aquatic and terrestrial settings. Nevertheless, paleoenvironmental data derived from these molecules have to be interpreted with care, as (i) their source microorganisms remain unknown, although some might belong to the phylum *Acidobacteria* (Sinninghe Damsté et al., 2011, 2018) and (ii) high uncertainty is associated with global brGDGT-mean annual air temperature (MAAT) calibrations (> 4 °C; Véquaud et al., 2022). The development of new environmental proxies, independent and complementary to brGDGTs, is crucial to improve the reliability and accuracy of continental reconstructions.

Recently, Wang et al. (2016) suggested that other bacterial lipids, so called 3-hydroxy fatty acids (3-OH FAs), could be used as such a proxy. These compounds, containing 10 to 18 C and a hydroxyl group in third position, are characteristic components of the Gram-negative bacterial membrane (e.g. Szponar et al., 2002, 2003; Keinänen et al., 2003). The membrane component of the cell wall of Gram-negative bacteria is more complex than that of Gram-positive cells, since it is composed of two distinct membranes, the outer membrane and the inner membrane, enclosing a thin space, the periplasm. The inner leaflet of the outer membrane is composed of phospholipids, while its outer leaflet is mainly composed of lipopolysaccharide (LPS) (Kamio and Nikaido, 1976). The LPS has a tripartite structure with the hydrophobic lipid A, the core oligosaccharide and the O-antigen polysaccharide, the two latter being hydrophilic. The lipid A, mainly constituted of 3-OH FAs (Raetz et al., 2007), anchors the LPS to the outer membrane and acts as a signaling molecule for the innate immune system. Three types of 3-OH FAs can be distinguished according to the presence and position of a methyl group on the carbon chain: *normal* (i.e. straight and unbranched), *iso* (methyl in penultimate position) or *anteiso* (methyl group in the antepenultimate position).

Gram-negative bacteria are ubiquitous in aquatic and terrestrial environments. 3-OH FAs were previously used to detect and quantify Gram-negative communities in various types of samples originating from terrestrial, aquatic and atmospheric environments (Lee et al., 2004; Wakeham et al., 2003; Tyagi et al., 2015, 2016). Nevertheless, until recently, these lipids have been largely overlooked for paleoclimate applications. Indeed, only limited information is available regarding the response of Gram-negative bacteria and associated 3-OH FA membrane lipids to the variations of environmental parameters.

Significant local correlations were recently obtained between the relative abundance of 3-OH FAs and temperature/pH in 26 soils collected along Mt. Shennongjia (China; Wang et al., 2016). Thus, the ratio of the summed *iso* and *anteiso* to the total amount of *normal* 3-OH FAs (the so-called RIAN index) was shown to vary with soil pH (Wang et al., 2016). In addition, the *anteiso* to *normal* 3-OH FA ratios of the C_15_ and C_17_ compounds (RAN_15_ and RAN_17_ indices, respectively) were negatively and linearly correlated with MAAT. The application of 3-OH FA based-proxies to a Holocene Chinese stalagmite reinforced the potential of these compounds as terrestrial paleoenvironmental proxies (Wang et al., 2018). Very recently, the applicability of 3-OH FAs as temperature and pH proxies was further investigated at the global level using extended soil datasets with samples from all over the world (Véquaud et al., 2021a; Wang et al., 2021). Significant relationships between the relative abundance of 3-OH FAs and temperature/pH at the global scale could only be obtained using non-linear models based on machine-learning algorithms (Véquaud et al., 2021a; Wang et al., 2021), highlighting the complexity of the response of 3-OH membrane lipids to changes in environmental parameters. Such a complexity may be related to the fact that (i) the diversity of Gram-negative bacteria varies from one soil to another (Margesin et al., 2009; Siles and Margesin, 2016) and that (ii) the 3-OH FA distribution differs between Gram-negative genera and species (Oyaizu and Komagata, 1983; Goossens et al., 1986; Hedrick et al., 2009; Wang et al., 2018). Therefore, the applicability of 3-OH FAs as environmental proxies in soils could be strongly influenced by the Gram-negative bacterial diversity, as the response to temperature and pH changes may vary between species. Nevertheless, to the best of our knowledge, no work to date has been performed at the microbial level to assess the impact of environmental parameters on the 3-OH FA profiles of various Gram-negative bacterial strains from soils.

In this study, we examined for the first time the influence of temperature and pH on three different strains of soil Gram-negative bacteria belonging to the same phylum (*Bacteroidetes*) in order to better understand the effect of variable environmental parameters on the membrane lipid profile, with a focus on 3-OH-FAs and their applicability as temperature and pH proxies. We hypothesized that (i) different temperature and pH culture conditions could lead to changes in 3-OH FA distribution and that (ii) the variations in 3-OH FA lipid profile in response to temperature and pH variations could differ from one strain to another.

## 2 Material and methods

### 2.1 Strain isolation

#### 2.1.1 Isolation of Gram-negative bacteria from surface soils

Gram-negative bacterial strains were isolated from surface soils (0–10 cm depth) collected in October 2017 along a well-documented composite altitudinal transect (234-2748 m) representative of the variations in temperature, soil characteristics and plant communities in the French Alps (Véquaud et al., 2021b). Isolation of Gram-negative bacterial strains was carried out from four contrasting soils collected in lowland, subalpine, mountainous and alpine land sites (samples 4, 19, 39 and 44, respectively, in Véquaud et al. (2021b). The physicochemical characteristics of these soil samples are detailed in Véquaud et al. (2021b).

Bacterial strains were isolated by resuspending two grams of each soil in 200 mL of sterile 0.9% (w/v) NaCl, incubated at 225 rpm on a rotary shaker for 20 min then left to sediment at room temperature for 40 min. The supernatant was diluted to 10^−3^ and 10^−4^ in sterile NaCl 0.9% before plating on R2A agar. Bacterial isolation was carried out at pH 7.2 or pH 5.0 (representative of the pH measured in the French Alps; Véquaud et al., 2021b) on R2A agar plates supplemented with vancomycin 10 mg/L, teicoplanin 10 mg/L, daptomycine 10 mg/L and cycloheximide 400 mg/L. For each soil, growth was monitored on eight plates at pH 7.2 and eight plates at pH 5.0, incubated at 5 °C or at 25 °C until no new colonies appeared. Twenty different clones covering the visible morphological differences of the colonies (colony size, shape, color, smoothness, general aspect), observed in each condition were re-isolated twice on the same medium. A single colony of each clone was then used to inoculate 10 mL of LB medium 0.1X, grown overnight at 25 °C or for 48h at 5 °C before storage of the cell pellet at −80 °C after resuspension in 60% glycerol, 7.5mM MgSO_4_. Three hundred and twenty cultivable species of Gram-negative bacteria were isolated from the four aforementioned soils (Véquaud et al., 2021b) and constituted the library for our study.

#### 2.1.2 Taxonomic identification

For each bacterial clone, a single colony was used for PCR amplification of rDNA sequences. Amplification was carried out with primers 27F and 1391R (Lane, 1991) using GoTaq® DNA polymerase (Promega) according to the manufacturer’s instructions, i.e. 95 °C for 2min, followed by 94 °C for 30s, 60 °C for 30 s, 72 °C for 1min40s (35 cycles), and 72 °C for 5 min. The PCR fragments were directly sequenced (Eurofins Genomics) using primer T7. A preliminary assignation was made using Standard Nucleotide Blast (Altschul et al., 1990). *Pseudomonas* generated a total of 193 first hits. The next most abundant genera were *Flavobacterium* (26 hits), *Colimonas* (17 hits), *Mucilaginibacter* (12 hits) and *Pedobacter* (9 hits). Three strains were selected from the library for further study, identified as belonging to the *Flavobacterium, Mucilaginibacter* and *Pedobacter* genera in the *Bacteroidetes* phylum. Further taxonomic affiliation was carried out on these clones after PCR amplification of the rDNA genes using the same primers as above and *Pfu* DNA polymerase (Promega), following the manufacturer’s instructions. The PCR fragments were cleaned, A-tailed and cloned into the pGEM-T vector. The identification of each strain was based on the sequencing of two positive plasmids using T7 and SP6 primers (Eurofins Genomics). The entire sequence was used for affiliation using EZBioCloud, database version 2121.07.07. The selected clones belonged to the *Bacteroidetes* phylum, as the C_15_ and C_17_ 3-OH FA homologues used in the RAN_15_ and RAN_17_ temperature proxies defined by Wang et al. (2016) were shown to be abundant in these bacteria (cf. section 4.1 for discussion).

The first strain, isolated from a calcosol collected at 232 m altitude (MAAT=12.5 °C; pH= 7.3; soil 44 in Véquaud et al., 2021b), was most closely related to *Flavobacterium pectinovorum* (98.78% similarity; GenBank accession no. OP204417). The second strain, isolated from a brunisol collected at 932 m altitude (MAAT=8.6 °C; pH=7.2; soil 39 in Véquaud et al., 2021b), was most closely related to *Pedobacter lusitanus* NL19 (98.54% similarity; GenBank accession no. OP204418). The third strain, isolated from a rendisol collected at 2695 m altitude (MAAT=0.4 °C; pH=6.5; soil 4 in Véquaud et al., 2021b), was most closely related to phylotype LGEL_s strain 048, *Pedobacter panaciterrae* (99.27% similarity; GenBank accession no. OP204416). In the rest of this paper, the three aforementioned strains will be referred to as strains A, B and C, respectively.

### 2.2 Cultivation of the strains

#### 2.2.1 Identification of the optimal growth conditions

For each strain, the optimal growth temperature at the intrinsic pH of the R2A medium (pH 7.2) was first identified. Triplicate cultures were performed for each temperature tested (5 °C, 10 °C, 15 °C, 20 °C, 25 °C and 30 °C). For each culture, a colony isolated on R2A agar medium was incubated in 7 mL of sterile R2A medium at 180 rpm (Infors HT Celltron) at the temperature to be tested to set up a subculture. After a few hours or a few days (depending on the strain and the tested temperature), the optical density (OD) of the subculture was measured at 600 nm (Spectrostarnano, BMG LabTech). One hundred ml of sterile R2A medium were inoculated with a volume of the subculture, chosen to reach an OD value between 0.05 and 0.10. The growth kinetics of each strain was then established. Cultures were shaken at 180 rpm under the desired conditions in a thermostatically controlled cabinet (Pol-Eko Aparatura®). OD was measured every hour until the stationary phase of bacterial growth was reached. The optimum pH (5, 6, 7 or 8) was defined under the same conditions, but with the temperature set at the defined optimal growth temperature (25 °C, see below). The pH of the R2A medium was adjusted with a pH meter using 1 M NaOH (pH above 7.2) or 1 M HCl (pH below 7.2). For each culture, bacterial growth curves were generated and the generation time was calculated in order to identify the optimal growth temperature and pH. 25 °C was identified as the optimal growth temperature for all the strains. At 25 °C, the optimal growth pH was 8 for strain A and 5 for strains B and C.

#### 2.2.2 Cultivation of the strains under different temperature and pH conditions

Once the optimum temperature and pH were identified, the strains were cultivated in triplicate in order to obtain a reference bacterial growth curve for each condition tested. The strains were cultivated at 180 rpm, either at fixed temperature (optimal growth temperature) and different pH (5, 6, 7 or 8) or at fixed pH (optimal growth pH) and different temperatures (5, 10, 15, 20 and 25 °C). These reference curves were used for regular measurements of OD, in order to determine the time until stationary phase is reached. Growth phase has been shown to have an influence on the membrane lipid composition of microorganisms (Männistö and Puhakka, 2001; Elling et al., 2014). Thus, in order to reliably compare the influence of temperature and pH on the distribution of 3-OH FAs between the different strains, the bacterial biomass was collected at the same stage of growth. For all the strains, the biomass was collected during the exponential phase by dividing by two the OD obtained at the stationary phase (reference OD). After subculturing as described above, the final cultures were performed in triplicate in a final volume of 150 mL of sterile R2A medium incubated at 180 rpm until reaching the reference OD. The cells were collected in 50 mL Falcon® tubes and centrifuged at 7000 rpm (JA12 rotor, Beckman) for 10 min at 20 °C. Each bacterial pellet was resuspended in 10 mL of 0.9 g/L NaCl to remove traces of R2A medium and then centrifuged again at the same speed as before. The supernatant was removed and the cell pellets were frozen at −20 °C before being freeze-dried for 48 h at −80 °C at a pressure of 0.93 Pa. The freeze-dried cell pellets were kept frozen at −20 °C before lipid extraction and analysis.

### 2.3 Lipid extraction

The freeze-dried cells were first subjected to acid methanolysis in 1M HCl /MeOH at 100 °C for 3 h. The suspension was then centrifuged at 15 °C and 3000 rpm for 5 min. The supernatant was collected in an Erlenmeyer flask. The pellet was extracted 3 times with a mixture of dichloromethane (DCM):MeOH (1:1, v/v; 25 mL). Each extraction was followed by centrifugation and pooling of all extracts. The extracts were then transferred to a separation funnel and washed with ultrapure water in order to neutralize the organic phase. The organic phase was recovered in a new Erlenmeyer flask. The aqueous phase was extracted twice with DCM (20 mL). The organic phase was recovered and pooled with the organic phase previously obtained. The organic phase containing the lipids was dried with sulfate sodium, then rotary-evaporated, diluted with a mixture of DCM:MeOH (5:1), transferred in a 4 mL vial and dried under nitrogen flow. The lipid extract was resuspended in 1 mL DCM and stored at −20 °C until 3-OH FA analysis.

### 2.4 Lipid analysis

The lipid extracts were first derivatised with a solution of *N,O-* bis(trimethylsilyl)trifluoroacetamide (BSTFA) at 70 °C for 45 min. Fatty acids were analysed by gas chromatography coupled to mass spectrometry using an Agilent Network 6980 GC System coupled with a 5973 Mass Selective Detector, with electron impact at 70 eV. A Restek RXI-5 Sil MS silica column (60 m x 0.25 mm, i.d. 0.50 μm film thickness) was used with He as the carrier gas at 1 mL/min, as previously described (Huguet et al., 2019). The GC oven program was: 70 °C to 200 °C at 10 °C/min, then to 310 °C (held 20min) at 2 °C/min. Samples were injected in splitless mode and the injector was set at 280 °C. The individual fatty acids (either hydroxy or non-hydroxy homologues) were first identified from the total ion chromatogram. The corresponding peaks were integrated, allowing the determination of the relative abundances of the different fatty acids.

A focus was then made on the different 3-OH FAs after extraction of the characteristic *m/z* 175 fragment (cf. Huguet et al., 2019), allowing the identification of the low-abundant homologues based on their relative retention times. 3-OH FAs were quantified by integrating the appropriate peak on the ion chromatogram and comparing the area with an analogous deuterated internal standard (3- hydroxytetradecanoic acid, 2,2,3,4,4-d5; Sigma-Aldrich, France). The internal standard (0.5 mg/mL) was added just before injection, with a proportion of 3 μL of standard to 100 μL of sample, as detailed by Huguet et al. (2019). The *m/z* 178 fragment was used for identification and integration of the deuterated internal standard.

The RIAN index was calculated as follows in the range C_10_-C_18_ (Wang et al., 2016):

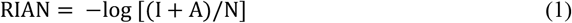

where I, A, N represent the sum of all *iso*, *anteiso* and *normal* 3-OH FAs, respectively. RAN_15_ and RAN_17_ indices were calculated as follows (Wang et al., 2016):

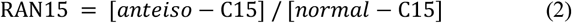

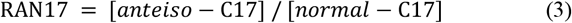

### 2.5 Statistical analyses

All the statistical analyses were performed using R (v3.6.3, Core Team, 2020). To investigate the statistical differences in the relative abundances of 3-OH FAs depending on the pH or temperature of the culture experiments, analysis of variance (ANOVA) and post-hoc Tukey tests were used. In order to investigate the correlations between the temperature or pH and the relative abundances of 3-OH FAs, pairwise Spearman correlation matrices and single linear regressions were performed for each strain. The *p*-value was set at 0.05 for all the statistical analyses.

## 3 Results

### 3.1 Global distribution of fatty acids in bacterial strains isolated from soils

#### 3.1.1 Non-hydroxy fatty acids

The membrane lipid profile of the three strains was investigated under different conditions of temperature and pH (Tables 1-3). The non-hydroxy FAs were generally more abundant than 3-OH FAs, especially for strain A, where non-hydroxy FAs represented more than 70 % of the total relative abundance in FAs (FAs; Table 1). For strains B and C, non-hydroxy FAs were generally slightly more abundant (> 55 % of total FAs) than 3-OH FAs. Straight-chain (i.e. *normal*) monounsaturated FAs were the major group of non-hydroxy FAs in strains B and C (Tables 2 and 3). In contrast, in strain A, the branched saturated FAs were as abundant as the normal monounsaturated FAs (Table 1). Strain A also differed from strains B and C by the presence of branched unsaturated FAs (*iso*-C_15:1_ and to a lesser extent *iso*-C_16:1_; Table 1). At the individual level, *normal*-C_16:1_ was observed to be the dominant non-hydroxy FA in strains B and C, followed, to a lesser extent, by *normal*-C_16:0_, *normal*-C_18:0_ and *iso*-C_15:0_ (Tables 2 and 3). Regarding strain A, the most abundant non-hydroxy FAs were the *iso*-C_15:0_ and *normal*-C_16:1_ homologues, followed by the *normal*-C_15:0_ and -C_16:0_ isomers.

**Table 1.** Relative abundances of non-hydroxy fatty acids (%) vs. total fatty acids for strain A at different temperature conditions (pH 8) and at different pH conditions (temperature 25 °C). Data are means of three biological replicates. Values after (±) are standard deviations. Correlations of the relative abundances of each fatty acid (FA) with either temperature or pH are provided with corresponding *p*-values. Significant correlations (*p* < 0.005) are indicated in bold. n.d.: not detected.

| Strain A | Temperature conditions (°C) |  |  |  |  |  |  | pH conditions |  |  |  |  |  |
| --- | --- | --- | --- | --- | --- | --- | --- | --- | --- | --- | --- | --- | --- |
|  | 5 °C | 10 °C | 15 °C | 20 °C | 25 °C | R<sup>2</sup> | *p* | 5 | 6 | 7 | 8 | R<sup>2</sup> | *p* |
| Normal saturated |  |  |  |  |  |  |  |  |  |  |  |  |  |
| 15:0 | 3.08 ± 0.16 | 4.60 ± 0.18 | 5.59 ± 0.42 | 7.45 ± 0.57 | 9.15 ± 1.10 | **0.94** | 2.11E-09 | 8.41 ± 3.27 | 12.39 ± 0.53 | 10.34 ± 2.34 | 9.15 ± 1.10 | 8.14E-05 | 0.98 |
| 16:0 | 8.75 ± 5.91 | 6.25 ± 1.29 | 6.13 ± 0.92 | 8.43 ± 0.74 | 12.81 ± 2.84 | 0.17 | 0.12 | 7.93 ± 2.36 | 7.50 ± 0.41 | 7.76 ± 0.70 | 12.81 ± 2.84 | **0.38** | 0.032 |
| 18:0 | 2.39 ± 0.83 | 3.71 ± 0.46 | 1.57 ± 0.76 | 1.27 ± 0.49 | 2.84 ± 0.42 | 0.05 | 0.45 | 0.86 ± 0.16 | 0.75 ± 0.28 | 0.62 ± 0.07 | 2.84 ± 0.42 | **0.48** | 0.013 |
| Total | 14.21 ± 5.83 | 14.57 ± 1.78 | 13.30 ± 1.50 | 17.17 ± 0.77 | 24.80 ± 4.11 | **0.45** | 6.29E-03 | 17.21 ± 5.44 | 20.62 ± 0.33 | 18.73 ± 2.95 | 24.80 ± 4.11 | 0.31 | 0.06 |
| Branched saturated |  |  |  |  |  |  |  |  |  |  |  |  |  |
| iso 14:0 | 0.63 ± 0.33 | 0.47 ± 0.03 | 0.19 ± 0.32 | n.d. | 0.68 ± 0.02 | 0.05 | 0.42 | 0.77 ± 0.26 | 0.42 ± 0.03 | 0.47 ± 0.05 | 0.68 ± 0.02 | 0.02 | 0.65 |
| iso 15:0 | 18.34 ± 0.37 | 19.27 ± 0.81 | 22.86 ± 3.19 | 24.18 ± 0.55 | 22.77 ± 1.65 | **0.54** | 1.75E-03 | 26.47 ± 0.93 | 25.37 ± 1.24 | 30.48 ± 1.16 | 22.77 ± 1.65 | 0.05 | 0.48 |
| anteiso 15:0 | 5.89 ± 0.07 | 4.59 ± 0.25 | 4.01 ± 0.43 | 2.67 ± 0.14 | 2.20 ± 0.20 | **0.96** | 3.88E-10 | 1.07 ± 0.11 | 2.06 ± 1.72 | 1.54 ± 0.15 | 2.20 ± 0.20 | 0.15 | 0.22 |
| iso 16:0 | 1.07 ± 0.31 | 1.48 ± 0.19 | 1.19 ± 0.21 | 0.86 ± 0.75 | 0.98 ± 0.03 | 0.06 | 0.37 | 0.09 ± 0.16 | n.d. | n.d. | 0.98 ± 0.03 | n.d. | n.d. |
| Total | 25.94 ± 0.73 | 25.82 ± 1.22 | 28.24 ± 3.11 | 27.71 ± 1.08 | 26.63 ± 1.85 | 0.07 | 0.34 | 28.40 ± 1.19 | 27.85 ± 1.10 | 32.50 ± 1.27 | 26.63 ± 1.85 | 8.33E-04 | 0.93 |
| Normal unsaturated |  |  |  |  |  |  |  |  |  |  |  |  |  |
| 15:1 | 1.41 ± 0.19 | 1.93 ± 0.18 | 1.92 ± 0.11 | 1.90 ± 0.09 | 1.86 ± 0.25 | 0.25 | 0.057 | 3.77 ± 0.70 | 2.70 ± 2.09 | 2.31 ± 0.41 | 1.86 ± 0.25 | **0.35** | 0.045 |
| 16:1 | 15.61 ± 0.35 | 15.58 ± 0.70 | 17.85 ± 1.34 | 17.88 ± 1.13 | 18.08 ± 0.63 | **0.58** | 1.05E-03 | 16.02 ± 2.56 | 15.49 ± 0.80 | 18.85 ± 0.87 | 18.08 ± 0.63 | **0.34** | 0.047 |
| 17:1 | 7.47 ± 2.93 | 10.86 ± 0.91 | 8.93 ± 0.81 | 4.95 ± 2.85 | 3.87 ± 1.81 | **0.37** | 0.017 | 4.00 ± 2.20 | 4.27 ± 0.87 | 3.15 ± 0.64 | 3.87 ± 1.81 | 0.017 | 0.69 |
| 18:1 | n.d. | n.d. | n.d. | n.d. | n.d. | n.d. | n.d. | n.d. | n.d. | n.d. | n.d. | n.d. | n.d. |
| Total | 24.49 ± 3.29 | 28.37 ± 0.75 | 28.45 ± 2.23 | 24.74 ± 2.45 | 23.81 ± 1.10 | 0.07 | 0.34 | 23.80 ± 1.59 | 22.46 ± 2.10 | 24.32 ± 1.89 | 23.81 ± 1.10 | 0.019 | 0.67 |
| Branched unsaturated |  |  |  |  |  |  |  |  |  |  |  |  |  |
| iso 15:1 | 9.73 ± 0.43 | 8.16 ± 0.44 | 6.96 ± 0.67 | 4.54 ± 0.19 | 4.04 ± 0.14 | **0.95** | 8.90E-10 | 3.41 ± 0.38 | 3.23 ± 0.14 | 3.39 ± 0.23 | 4.04 ± 0.14 | **0.38** | 0.032 |
| iso 16:1 | 0.22 ± 0.38 | 0.48 ± 0.42 | 0.24 ± 0.41 | n.d. | n.d. | **0.56** | 1.27E-03 | n.d. | n.d. | n.d. | n.d. | 0.09 | 0.14 |
| Total | 9.95 ± 0.54 | 8.64 ± 0.36 | 7.20 ± 0.33 | 4.54 ± 0.19 | 4.04 ± 0.14 | **0.95** | 4.29E-10 | 3.41 ± 0.38 | 3.23 ± 0.14 | 3.39 ± 0.23 | 4.04 ± 0.14 | **0.38** | 0.03 |
| Total non-hydroxy FAs | 74.60 ± 2.86 | 77.40 ± 2.90 | 77.18 ± 5.06 | 74.15 ± 3.22 | 79.28 ± 5.00 | 0.05 | 0.41 | 72.81 ± 5.95 | 74.16 ± 1.55 | 78.93 ± 5.83 | 79.28 ± 5.00 | 0.30 | 0.065 |

**Table 2.** Relative abundances of non-hydroxy fatty acids (%) vs. total fatty acids for strain B at different temperature conditions (pH 5) and at different pH conditions (temperature 25 °C). Data are means of three biological replicates. Values after (±) are standard deviations. Correlations of the relative abundances of each fatty acid (FA) with either temperature or pH are provided with corresponding *p*-values. Significant correlations *(p* < 0.005) are indicated in bold. n.d.: not detected.

| Strain B | Temperature conditions (°C) |  |  |  |  |  |  | pH conditions |  |  |  |  |  |
| --- | --- | --- | --- | --- | --- | --- | --- | --- | --- | --- | --- | --- | --- |
|  | 5 °C | 10 °C | 15 °C | 20 °C | 25 °C | R<sup>2</sup> | *p* | 5 | 6 | 7 | 8 | R<sup>2</sup> | *p* |
| **Normal saturated** |  |  |  |  |  |  |  |  |  |  |  |  |  |
| 15:0 | n.d. | n.d. | n.d. | n.d. | n.d. | n.d. | n.d. | n.d. | n.d. | n.d. | n.d. | n.d. | n.d. |
| 16:0 | 11.80 ± 0.99 | 13.18 ± 3.99 | 12.13 ± 0.98 | 11.31 ± 1.59 | 12.09 ± 1.44 | 0.010 | 0.73 | 12.09 ± 1.44 | 13.94 ± 4.77 | 12.69 ± 1.02 | 17.08 ± 1.84 | **0.27** | 0.081 |
| 18:0 | 4.94 ± 1.54 | 5.90 ± 3.17 | 3.42 ± 0.41 | 6.19 ± 3.21 | 2.39 ± 0.62 | **0.23** | 0.28 | 2.39 ± 0.62 | 1.87 ± 0.87 | 2.40 ± 0.64 | 3.62 ± 0.67 | 0.30 | 0.067 |
| Total | 16.74 ± 2.53 | 19.07 ± 5.50 | 15.56 ± 1.39 | 17.50 ± 4.80 | 14.48 ± 2.06 | 0.83 | 0.35 | 14.48 ± 2.06 | 15.82 ± 3.91 | 15.08 ± 1.35 | 20.70 ± 2.28 | 0.38 | 0.032 |
| **Branched saturated** |  |  |  |  |  |  |  |  |  |  |  |  |  |
| iso 14:0 | n.d. | 2.80 ± 2.52 | n.d. | n.d. | 0.13 ± 0.22 | n.d. | n.d. | 0.13 ± 0.22 | 0.31 ± 0.53 | 0.22 ± 0.38 | n.d. | 0.26 | 0.092 |
| iso 15:0 | 5.61 ± 0.37 | 7.09 ± 1.08 | 6.03 ± 0.32 | 7.27 ± 0.61 | 10.10 ± 1.00 | **0.59** | 7.83E-04 | 10.10 ± 1.00 | 6.35 ± 0.99 | 6.50 ± 1.22 | 9.23 ± 2.24 | 0.02 | 0.68 |
| anteiso 15:0 | 2.52 ± 0.22 | 1.72 ± 1.54 | 1.49 ± 0.16 | 0.25 ± 0.44 | 0.23 ± 0.40 | **0.81** | 4.84E-06 | 0.23 ± 0.40 | n.d. | 0.14 ± 0.24 | n.d. | **n.d.** | 1.30E-02 |
| iso 16:0 | n.d. | n.d. | n.d. | n.d. | n.d. | n.d. | n.d. | n.d. | n.d. | n.d. | n.d. | n.d. | n.d. |
| Total | 8.13 ± 0.59 | 11.60 ± 1.37 | 7.52 ± 0.20 | 7.53 ± 0.73 | 10.45 ± 1.62 | 0.002 | 0.88 | 10.45 ± 1.62 | 6.65 ± 0.48 | 6.85 ± 0.91 | 9.23 ± 2.24 | 0.04 | 0.55 |
| **Normal unsaturated** |  |  |  |  |  |  |  |  |  |  |  |  |  |
| 15:1 | n.d. | n.d. | n.d. | n.d. | n.d. | n.d. | n.d. | n.d. | n.d. | n.d. | n.d. | n.d. | n.d. |
| 16:1 | 33.69 ± 1.43 | 33.71 ± 1.87 | 32.65 ± 2.33 | 33.11 ± 2.59 | 30.86 ± 1.29 | 0.21 | 0.083 | 30.86 ± 1.29 | 33.05 ± 1.30 | 31.54 ± 2.03 | 7.70 ± 0.78 | **0.57** | 4.68E-03 |
| 17:1 | 7.89 ± 1.70 | 4.49 ± 1.41 | 2.33 ± 0.28 | n.d. | 0.57 ± 0.52 | **0.77** | 1.85E-05 | 0.57 ± 0.52 | 0.46 ± 0.09 | 0.71 ± 0.05 | n.d. | **0.05** | 0.49 |
| 18:1 | 0.36 ± 0.34 | n.d. | n.d. | n.d. | 0.32 ± 0.29 | 0.09 | 0.29 | 0.32 ± 0.29 | 3.18 ± 5.50 | n.d. | n.d. | n.d. | n.d. |
| Total | 41.94 ± 0.38 | 38.20 ± 0.78 | 34.99 ± 2.04 | 33.11 ± 2.59 | 31.76 ± 2.05 | **0.83** | 2.31E-06 | 31.76 ± 2.05 | 36.69 ± 6.62 | 32.25 ± 1.99 | 7.70 ± 0.78 | 0.53 | 7.10E-03 |
| **Branched unsaturated** |  |  |  |  |  |  |  |  |  |  |  |  |  |
| iso 15:1 | n.d. | n.d. | n.d. | n.d. | n.d. | n.d. | n.d. | n.d. | n.d. | n.d. | n.d. | n.d. | n.d. |
| Iso 16:1 | n.d. | n.d. | n.d. | n.d. | n.d. | n.d. | n.d. | n.d. | n.d. | n.d. | n.d. | n.d. | n.d. |
| Total | n.d. | n.d. | n.d. | n.d. | n.d. | n.d. | n.d. | n.d. | n.d. | n.d. | n.d. | n.d. | n.d. |
| **Total non-hydroxy FAs** | **66.82 ± 2.89** | **68.89 ± 6.23** | **58.06 ± 3.62** | **58.13 ± 6.64** | **56.69 ± 5.08** | **0.45** | 6.54E-03 | **56.69 ± 5.08** | **59.16 ± 10.09** | **54.18 ± 3.91** | **37.62 ± 0.10** | **0.51** | 9.20E-03 |

**Table 3.**
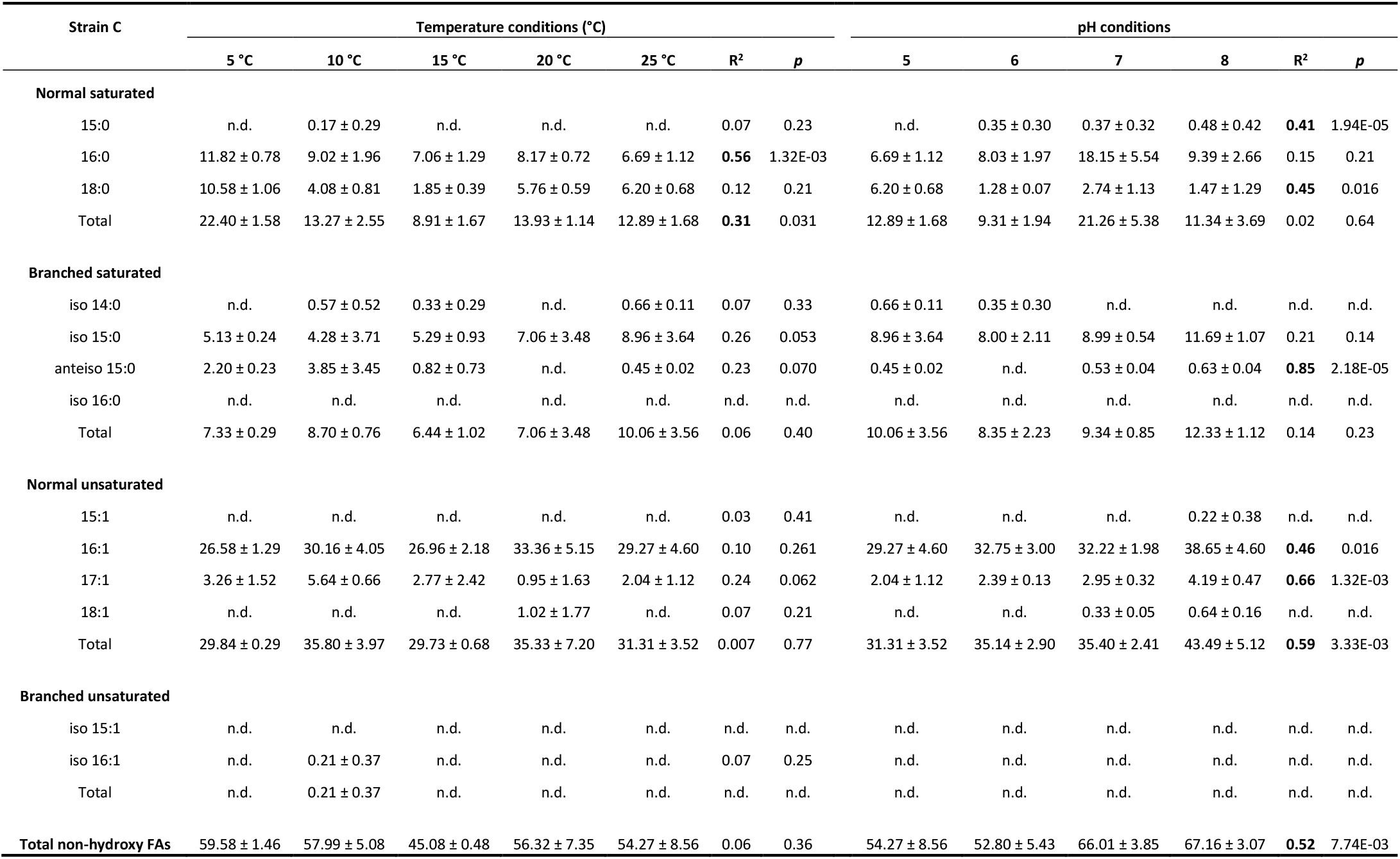
Relative abundances of non-hydroxy fatty acids (%) vs. total fatty acids for strain C at different temperature conditions (pH 5) and at different pH conditions (temperature 25 °C). Data are means of three biological replicates. Values after (±) are standard deviations. Correlations of the relative abundances of each fatty acid (FA) with either temperature or pH are provided with corresponding *p*-values. Significant correlations (*p* < 0.005) are indicated in bold. n.d.: not detected.

#### 3.1.2 3-OH FAs

The distribution of the 3-OH FAs in the three investigated strains was also examined in detail (Figure 1 and Tables 4-6). The 3-OH FAs detected had a carbon number of between 13C and 18C. Whatever the cultivation conditions, the *iso* homologues were the most abundant for strain A (Table 4). This was also the case for strain B, except at a growth temperature of 5 °C (Table 5), where the *anteiso* homologues were slightly more abundant than the *iso* homologues. In addition, the relative abundance of the *normal* homologues was higher than the one of the *anteiso* isomers for strain A (Table 4), whereas an opposite trend was observed for strain C (Table 6). Regarding strain B, the relative abundance of *normal* vs. *anteiso* isomers was highly dependent on the cultivation conditions (Table 4).

**Figure 1:**
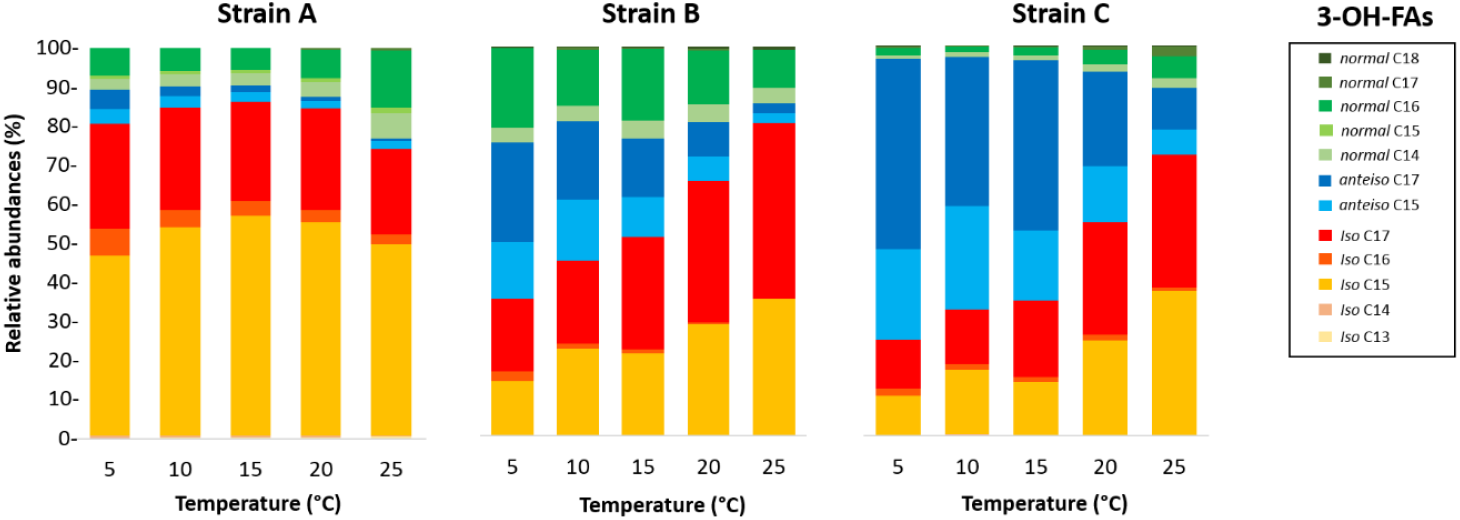
Variation of the relative abundances of 3-OH-FAs with temperature for the three Gram-negative bacterial strains isolated from soils of the French Alps. Strain A: *Flavobacterium pectinovorum* (98.78% similarity). Strain B: *Pedobacter lusitanus* NL19 (98.54% similarity). Strain C: *Pedobacter panaciterrae* (99.27% similarity)

**Table 4.** Relative abundances of individual 3-hydroxy fatty acids (%) vs. total 3-hydroxy fatty acids and related indices (defined by Wang et al., 2016) for strain A at different temperature conditions (pH 8) and at different pH conditions (temperature 25 °C). Data are means of three biological replicates. Values after (±) are standard deviations. Correlations of the relative abundances of each 3-hydroxy fatty acid with either temperature or pH are provided with corresponding *p*-values. Significant correlations (*p* < 0.005) are indicated in bold. n.d.: not detected.

| Strain A | Temperature conditions (°C) |  |  |  |  |  |  | pH conditions |  |  |  |  |  |
| --- | --- | --- | --- | --- | --- | --- | --- | --- | --- | --- | --- | --- | --- |
|  | 5 °C | 10 °C | 15 °C | 20 °C | 25 °C | R<sup>2</sup> | *p* | 5 | 6 | 7 | 8 | R<sup>2</sup> | *p* |
| iso C<sub>13</sub> | 0.15 ± 0.006 | 0.28 ± 0.02 | 0.33 ± 0.06 | 0.42 ± 0.03 | 0.48 ± 0.03 | **0.91** | 2.59E-08 | 1.15 ± 0.36 | 0.79 ± 0.08 | 0.54 ± 0.21 | 0.48 ± 0.03 | **0.64** | 1.73E-03 |
| iso C<sub>14</sub> | 0.52 ± 0.03 | 0.45 ± 0.04 | 0.44 ± 0.15 | 0.37 ± 0.04 | 0.31 ± 0.001 | **0.55** | 1.45E-03 | n.d. | 0.09 ± 0.08 | 0.21 ± 0.07 | 0.31 ± 0.001 | **0.88** | 7.93E-06 |
| normal C<sub>14</sub> | 2.88 ± 0.15 | 3.06 ± 0.45 | 3.22 ± 0.31 | 3.69 ± 0.08 | 6.35 ± 0.88 | **0.64** | 3.51E-04 | 7.81 ± 2.25 | 4.66 ± 0.52 | 5.03 ± 1.05 | 6.35 ± 0.88 | 0.07 | 0.39 |
| iso C<sub>15</sub> | 46.17 ± 1.43 | 53.41 ± 5.32 | 56.37 ± 10.02 | 54.64 ± 4.40 | 49.02 ± 6.17 | 0.02 | 0.58 | 65.92 ± 7.93 | 64.32 ± 4.78 | 59.69 ± 9.43 | 49.02 ± 6.17 | **0.48** | 0.012 |
| anteiso C<sub>15</sub> | 3.74 ± 0.16 | 3.08 ± 0.26 | 2.45 ± 0.29 | 1.88 ± 0.22 | 1.98 ± 0.19 | **0.85** | 1.10E-6 | 0.61 ± 0.11 | 0.71 ± 0.09 | 1.28 ± 0.34 | 1.98 ± 0.19 | **0.84** | 2.74E-05 |
| normal C<sub>15</sub> | 0.72 ± 0.008 | 0.79 ± 0.08 | 0.81 ± 0.28 | 1.10 ± 0.05 | 1.60 ± 0.08 | **0.73** | 4.78E-05 | 2.35 ± 0.32 | 2.41 ± 0.03 | 1.61 ± 0.28 | 1.60 ± 0.08 | **0.64** | 1.79E-03 |
| iso C<sub>16</sub> | 6.92 ± 0.62 | 4.27 ± 0.89 | 3.77 ± 0.28 | 3.14 ± 0.54 | 2.47 ± 0.68 | **0.68** | 1.42E-04 | 0.58 ± 0.35 | 0.71 ± 0.10 | 1.07 ± 0.32 | 2.47 ± 0.68 | **0.67** | 1.09E-03 |
| normal C<sub>16</sub> | 7.10 ± 0.21 | 5.88 ± 0.76 | 5.53 ± 2.43 | 7.44 ± 1.64 | 14.45 ± 2.69 | **0.41** | 9.94E-03 | 10.20 ± 1.90 | 8.94 ± 1.58 | 12.29 ± 5.48 | 14.45 ± 2.69 | 0.28 | 0.078 |
| iso C<sub>17</sub> | 26.93 ± 0.79 | 26.30 ± 3.79 | 25.40 ± 3.92 | 25.95 ± 2.23 | 21.89 ± 3.66 | 0.23 | 0.073 | 10.84 ± 7.68 | 16.33 ± 3.67 | 17.17 ± 5.92 | 21.89 ± 3.66 | **0.41** | 0.026 |
| anteiso C<sub>17</sub> | 4.81 ± 0.30 | 2.44 ± 0.53 | 1.62 ± 0.86 | 1.05 ± 0.19 | 0.75 ± 0.23 | **0.79** | 8.29E-06 | 0.13 ± 0.12 | 0.32 ± 0.05 | 0.56 ± 0.13 | 0.75 ± 0.23 | **0.79** | 1.07E-04 |
| normal C<sub>17</sub> | 0.07 ± 0.06 | 0.04 ± 0.07 | 0.07 ± 0.12 | 0.32 ± 0.04 | 0.70 ± 0.05 | **0.71** | 7.94E-05 | 0.42 ± 0.37 | 0.72 ± 0.13 | 0.54 ± 0.07 | 0.70 ± 0.05 | 0.14 | 0.23 |
| normal C<sub>18</sub> | n.d. | n.d. | n.d. | n.d. | n.d. | n.d. | n.d. | n.d. | n.d. | n.d. | n.d. | n.d. | n.d. |
| Total iso 3-OH FAs | 80.69 ± 0.16 | 84.71 ± 0.74 | 86.31 ± 4.27 | 84.52 ± 1.75 | 74.17 ± 1.88 | 0.16 | 0.15 | 78.49 ± 1.03 | 82.24 ± 1.18 | 78.69 ± 3.85 | 74.17 ± 1.88 | 0.29 | 0.070 |
| Total anteiso 3-OH FAs | 8.55 ± 0.13 | 5.52 ± 0.29 | 4.07 ± 1.15 | 2.93 ± 0.35 | 2.72 ± 0.05 | **0.85** | 1.15E-06 | 0.73 ± 0.02 | 1.04 ± 0.05 | 1.84 ± 0.35 | 2.72 ± 0.05 | **0.93** | 5.20E-07 |
| Total normal 3-OH FAs | 10.77 ± 0.13 | 9.77 ± 0.45 | 9.62 ± 3.13 | 12.56 ± 1.61 | 23.10 ± 1.86 | **0.54** | 1.81E-03 | 20.78 ± 1.04 | 16.72 ± 1.22 | 19.47 ± 4.20 | 23.10 ± 1.86 | 0.12 | 0.25 |
| Iso/anteiso 3-OH FAs | 9.44 ± 0.15 | 15.39 ± 0.97 | 22.42 ± 6.39 | 29.16 ± 3.75 | 27.27 ± 1.06 | **0.79** | 1.05E-05 | 107.0 ± 2.81 | 79.48 ± 3.30 | 43.57 ± 7.11 | 27.27 ± 1.06 | **0.97** | 6.22E-09 |
| Iso/normal 3-OH FAs | 7.50 ± 0.10 | 8.68 ± 0.49 | 9.65 ± 3.17 | 6.82 ± 1.06 | 3.23 ± 0.36 | **0.34** | 0.023 | 3.79 ± 0.23 | 4.94 ± 0.45 | 4.19 ± 1.06 | 3.23 ± 0.36 | 0.11 | 0.28 |
| Anteiso/normal 3-OH FAs | 0.79 ± 0.02 | 0.56 ± 0.005 | 0.43 ± 0.03 | 0.24 ± 0.04 | 0.12 ± 0.009 | **0.98** | 6.79E-13 | 0.04 ± 0.02 | 0.06 ± 0.008 | 0.10 ± 0.04 | 0.12 ± 0.009 | **0.79** | 1.05E-04 |
| RIAN | -0.92 ± 0.006 | -0.97 ± 0.02 | -0.99 ± 0.15 | -0.85 ± 0.07 | -0.52 ± 0.05 | **0.51** | 2.87E-03 | -0.58 ± 0.03 | -0.70 ± 0.04 | -0.62 ± 0.12 | -0.52 ± 0.05 | 0.11 | 0.29 |
| RAN<sub>15</sub> | 5.22 ± 0.27 | 3.96 ± 0.75 | 3.17 ± 0.67 | 1.70 ± 0.16 | 1.24 ± 0.14 | **0.92** | 2.31E-08 | 0.26 ± 0.07 | 0.30 ± 0.04 | 0.79 ± 0.08 | 1.24 ± 0.14 | **0.88** | 5.57E-06 |
| RAN<sub>17</sub> | 44.80 ± 2.33 | 24.44 | 12.51 | 3.30 ± 0.21 | 1.07 ± 0.34 | **0.92** | 1.26E-08 | 0.31 ± 0.03 | 0.46 ± 0.02 | 1.02 ± 0.10 | 1.07 ± 0.34 | **0.73** | 4.20E-04 |

**Table 5.**
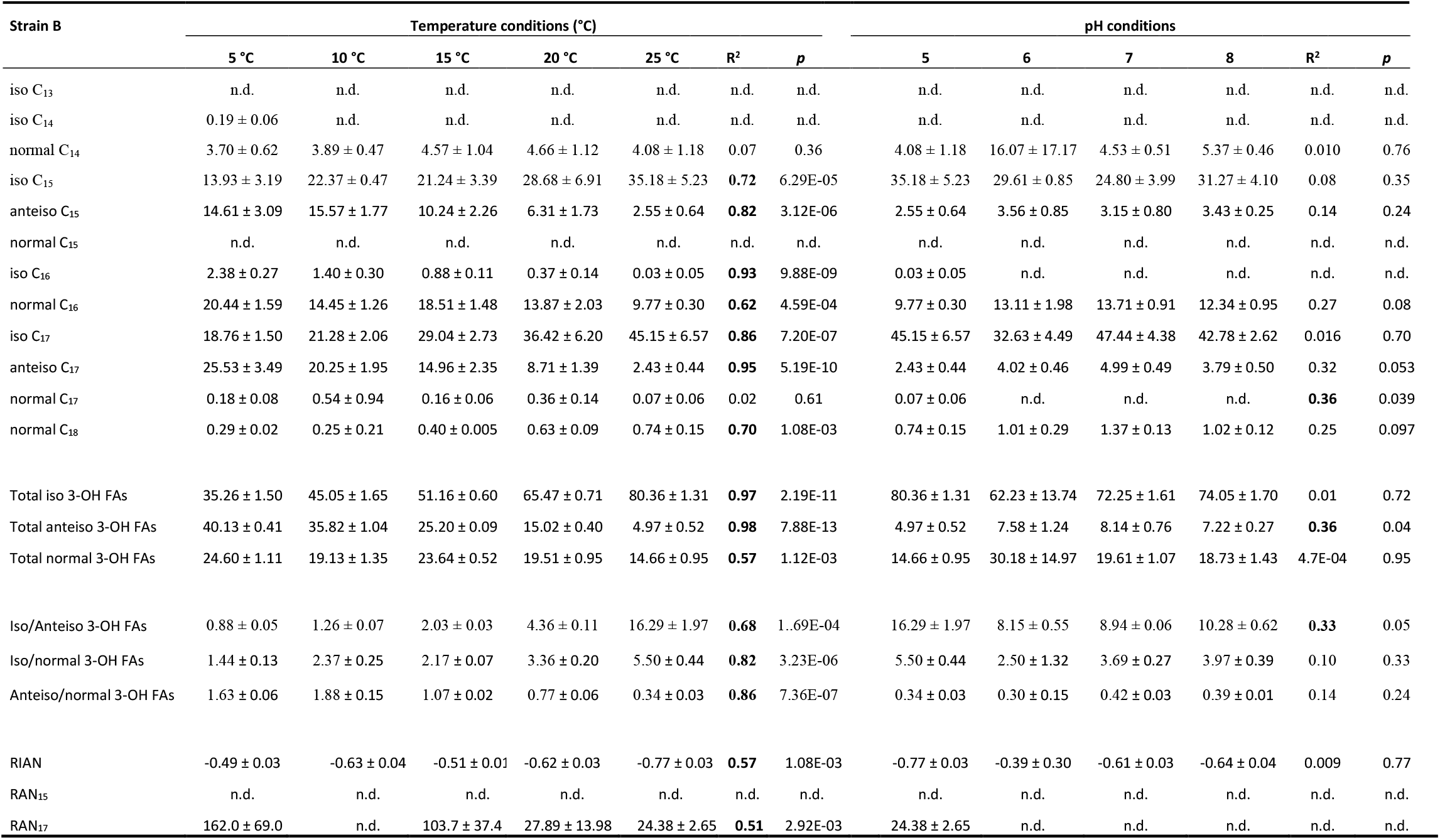
Relative abundances of individual 3-hydroxy fatty acids (%) vs. total 3-hydroxy fatty acids and related indices (defined by Wang et al., 2016) for strain B at different temperature conditions (pH 5) and at different pH conditions (temperature 25 °C). Data are means of three biological replicates. Values after (±) are standard deviations. Correlations of the relative abundances of each 3-hydroxy fatty acid with either temperature or pH are provided with corresponding *p*-values. Significant correlations (*p* < 0.005) are indicated in bold. n.d.: not detected.

**Table 6.** Relative abundances of individual 3-hydroxy fatty acids (%) vs. total 3-hydroxy fatty acids and related indices (defined by Wang et al., 2016) for strain C at different temperature conditions (pH 8) and at different pH conditions (temperature 25 °C). Data are means of three biological replicates. Values after (±) are standard deviations. Correlations of the relative abundances of each 3-hydroxy fatty acid with either temperature or pH are provided with corresponding *p*-values. Significant correlations (*p* < 0.005) are indicated in bold. n.d.: not detected.

| Strain C | Temperature conditions (°C) |  |  |  |  |  |  | pH conditions |  |  |  |  |  |
| --- | --- | --- | --- | --- | --- | --- | --- | --- | --- | --- | --- | --- | --- |
| | 5 °C | 10 °C | 15 °C | 20 °C | 25 °C | R<sup>2</sup> | $p$ | 5 | 6 | 7 | 8 | R<sup>2</sup> | $p$ |
| iso C<sub>13</sub> | n.d. | n.d. | n.d. | n.d. | n.d. | n.d. | n.d. | n.d. | n.d. | n.d. | n.d. | n.d. | n.d. |
| iso C<sub>14</sub> | 0.31 ± 0.04 | 0.34 ± 0.14 | 0.16 ± 0.01 | 0.10 ± 0.09 | 0.03 ± 0.06 | **0.69** | 1.39E-04 | 0.03 ± 0.06 | 0.12 ± 0.06 | 0.23 ± 0.06 | 0.27 ± 0.04 | **0.79** | 1.24E-04 |
| normal C<sub>14</sub> | 0.79 ± 0.17 | 1.25 ± 0.54 | 1.32 ± 0.33 | 2.01 ± 0.26 | 2.48 ± 1.40 | **0.50** | 3.37E-03 | 2.48 ± 1.40 | 3.55 ± 1.29 | 3.11 ± 0.64 | 3.17 ± 0.62 | 0.04 | 0.54 |
| iso C<sub>15</sub> | 9.98 ± 1.84 | 16.57 ± 4.20 | 13.60 ± 0.82 | 24.36 ± 5.12 | 37.08 ± 9.96 | **0.68** | 1.41E-04 | 37.08 ± 9.96 | 36.44 ± 5.78 | 22.08 ± 2.60 | 26.93 ± 4.94 | **0.37** | 0.036 |
| anteiso C<sub>15</sub> | 23.04 ± 5.28 | 26.57 ± 5.97 | 17.95 ± 1.64 | 14.45 ± 3.63 | 6.45 ± 3.19 | **0.67** | 1.73E-04 | 6.45 ± 3.19 | 6.33 ± 1.00 | 4.55 ± 0.49 | 6.16 ± 1.26 | 0.03 | 0.58 |
| normal C<sub>15</sub> | n.d. | n.d. | n.d. | n.d. | n.d. | n.d. | n.d. | n.d. | 0.14 ± 0.25 | 0.13 ± 0.02 | 0.17 ± 0.07 | 0.19 | 0.15 |
| iso C<sub>16</sub> | 1.78 ± 0.23 | 1.54 ± 0.32 | 1.28 ± 0.13 | 1.40 ± 0.39 | 0.91 ± 0.05 | **0.55** | 1.47E-03 | 0.91 ± 0.05 | 1.02 ± 0.40 | 2.29 ± 0.18 | 1.79 ± 0.34 | **0.51** | 8.88E-03 |
| normal C<sub>16</sub> | 1.91 ± 0.48 | 1.54 ± 0.49 | 2.09 ± 0.30 | 3.77 ± 1.64 | 5.58 ± 1.78 | **0.58** | 9.16E-04 | 5.58 ± 1.78 | 5.03 ± 1.13 | 8.26 ± 0.52 | 7.54 ± 1.41 | **0.36** | 0.040 |
| iso C<sub>17</sub> | 12.65 ± 2.68 | 13.85 ± 2.78 | 19.64 ± 0.83 | 28.86 ± 2.92 | 34.01 ± 7.05 | **0.83** | 2.03E-06 | 34.01 ± 7.05 | 37.15 ± 7.24 | 48.26 ± 1.93 | 43.61 ± 3.99 | **0.39** | 0.029 |
| anteiso C<sub>17</sub> | 48.96 ± 3.95 | 38.19 ± 8.68 | 43.64 ± 1.73 | 24.11 ± 4.32 | 10.81 ± 5.00 | **0.77** | 1.61E-05 | 10.81 ± 5.00 | 9.65 ± 1.40 | 10.21 ± 1.93 | 9.25 ± 1.13 | 0.04 | 0.55 |
| normal C<sub>17</sub> | 0.50 ± 0.26 | 0.14 ± 0.02 | 0.19 ± 0.08 | 0.76 ± 1.07 | 2.41 ± 1.25 | **0.37** | 0.016 | 2.41 ± 1.25 | 0.30 ± 0.27 | 0.48 ± 0.26 | 0.71 ± 0.07 | 0.31 | 0.062 |
| normal C<sub>18</sub> | 0.07 ± 0.06 | n.d. | 0.12 ± 0.03 | 0.18 ± 0.03 | 0.24 ± 0.10 | **0.59** | 8.08E-04 | 0.24 ± 0.10 | 0.27 ± 0.03 | 0.40 ± 0.05 | 0.41 ± 0.09 | **0.53** | 7.21E-03 |
| Total iso 3-OH FAs | 24.71 ± 1.84 | 32.30 ± 2.07 | 34.69 ± 0.80 | 54.72 ± 3.29 | 72.03 ± 3.52 | **0.90** | 5.75E-08 | 72.03 ± 3.52 | 74.73 ± 1.25 | 72.85 ± 2.42 | 72.60 ± 0.89 | 7.99E-5 | 0.98 |
| Total anteiso 3-OH FAs | 72.00 ± 2.27 | 64.76 ± 3.05 | 61.59 ± 1.00 | 38.56 ± 0.71 | 17.26 ± 7.54 | **0.88** | 1.97E-07 | 17.26 ± 7.54 | 15.97 ± 0.50 | 14.76 ± 2.25 | 15.41 ± 0.18 | 0.05 | 0.48 |
| Total normal 3-OH FAs | 3.28 ± 0.57 | 2.94 ± 0.98 | 3.72 ± 0.22 | 6.72 ± 2.75 | 10.71 ± 4.18 | **0.57** | 1.12E-03 | 10.71 ± 4.18 | 9.30 ± 1.75 | 12.38 ± 0.18 | 11.99 ± 0.93 | 0.12 | 0.27 |
| Iso/anteiso 3-OH FAs | 0.34 ± 0.04 | 0.50 ± 0.06 | 0.56 ± 0.02 | 1.42 ± 0.11 | 5.09 ± 3.17 | **0.48** | 4.41E-03 | 5.09 ± 3.17 | 4.68 ± 0.07 | 5.03 ± 0.91 | 4.71 ± 0.08 | 4.23E-3 | 0.84 |
| Iso/normal 3-OH FAs | 7.63 ± 0.97 | 11.61 ± 2.82 | 9.33 ± 0.37 | 9.07 ± 3.40 | 7.26 ± 2.08 | 0.04 | 0.49 | 7.26 ± 2.08 | 8.26 ± 1.76 | 5.89 ± 0.28 | 6.08 ± 0.55 | 0.20 | 0.15 |
| Anteiso/normal 3-OH FAs | 22.46 ± 4.64 | 23.80 ± 7.98 | 16.59 ± 1.20 | 6.29 ± 2.04 | 1.92 ± 1.18 | **0.78** | 1.33E-05 | 1.92 ± 1.18 | 1.77 ± 0.40 | 1.19 ± 0.16 | 1.29 ± 0.11 | 0.21 | 0.14 |
| RIAN | -1.47 ± 0.08 | -1.53 ± 0.14 | -1.41 ± 0.03 | -1.16 ± 0.18 | -0.94 ± 0.18 | **0.67** | 1.74E-04 | -0.94 ± 0.18 | -0.99 ± 0.09 | -0.85 ± 0.007 | -0.87 ± 0.04 | 0.16 | 0.19 |
| RAN<sub>15</sub> | n.d. | n.d. | n.d. | n.d. | n.d. | n.d. | n.d. | n.d. | 17.39 | 34.49 ± 7.10 | 40.13 ± 11.40 | **0.45** | 0.017 |
| RAN<sub>17</sub> | 126.5 ± 86.2 | 275.4 ± 91.9 | 258.3 ± 127.3 | 46.6 ± 45.6 | 6.27 ± 5.46 | 0.25 | 0.057 | 6.27 ± 5.46 | 22.98 ± 2.18 | 23.58 ± 7.50 | 13.10 ± 0.33 | 0.11 | 0.29 |

The distribution of the individual 3-OH FAs was also observed to differ between the strains. Thus, for strain A, the *iso*-C_15_ homologue was by far the most abundant 3-OH FA, followed by the *iso*-C_17_ and *normal*-C_16_ compounds, whatever the cultivation conditions (Figure 1 and Table 4). Regarding strain B, the 3-OH FA distribution was dominated by either *iso*-C_15_, *iso*-C_17_ or *anteiso*-C_17_ homologues (depending on the cultivation conditions), followed by the *n*-C_16_ isomer (Table 4). As for strain C, the most abundant 3-OH FAs were *anteiso*-C_17_, *iso*-C_17_, *anteiso*-C_15_ and *iso*-C_15_, with a strong effect of the cultivation conditions on the relative proportion of these isomers. It should be noted that part of the 3-OH FAs were present at very low abundance (< 1% of total abundance in 3-OH FAs), and were even not detected in some of the samples. For example, the *iso*-C_13_ and *normal*-C_15_ homologs were mainly detected in strain A isolates (Table 4). In contrast, the *normal*-C_18_ isomer was only detected in strains B and C (Tables 5 and 6).

### 3.2 Influence of temperature on the FA distribution of the different strains

#### 3.2.1 Non-hydroxy fatty acids

The influence of temperature on the relative abundance of the individual FAs was compound- and strain-dependent. Regarding the non-hydroxy fatty acids, the relative abundance of *anteiso*-C_15:0_ and *normal*-C_17:1_ was observed to significantly decrease with an increase in temperature for both strains A and B (Tables 1 and 2). In addition to the two aforementioned homologues, temperature variations had a strong influence on the relative abundance of other isomers in strain A, as the relative abundances in *normal*-C_15:0_ and *iso*-C_15:0_ were shown to significantly increase with temperature (R^2^ = 0.94 and 0.54, respectively), whereas an opposite trend was observed for *iso*-C_15:1_ (R^2^ = 0.95; Table 1). In contrast, temperature had only a limited impact on the non-hydroxy FA lipid profile in strain C (Table 3).

#### 3.2.2 3-OH FAs

In all strains, the relative abundance of the total *anteiso* 3-OH FAs was observed to significantly decrease with increasing temperature (Tables 4-6). In addition, in strains A and C, the relative abundance of the total *normal* 3-OH FAs was significantly and positively correlated with temperature (Tables 4 and 6), whereas an opposite trend was observed for strain B (Table 5). In strains B and C, the relative abundance of the total *iso* 3-OH FAs was also positively and strongly correlated with temperature (Table 5 and 6).

The relative abundance of some of the individual 3-OH FAs was also observed to be similarly affected by temperature changes, whatever the strain. Thus, in strain C, the relative abundance of the *anteiso*-C_17_ and *anteiso*-C_15_ isomers significantly decreased with temperature (R^2^ = 0.77 and 0.67, respectively; Figure 1 and Table 6). Such a significant decrease in the relative abundances of the *anteiso*-C_17_ and *anteiso*-C_15_ 3-OH FAs with temperature was similarly observed in strains B (R^2^ = 0.95 and 0.82, respectively; Table 5) and strain A (R^2^ = 0.79 and 0.85, respectively; Table 4) despite the low abundance of the two *anteiso* isomers in the latter strain (< 5% of total 3-OH FAs). It should be noted that the relative abundance of the *iso*-C_14_ 3-OH FA was also observed to significantly decrease with temperature in strains A and C (R^2^ = 0.55 and 0.69, respectively; Tables 4 and 6), despite its low proportion (this compound was even not detected in most of the strain B samples).

In contrast, the effect of temperature differed between the strains for some of the individual 3-OH FAs. The relative abundance of the *iso*-C_15_ isomer, predominant in strain A, did not change significantly with temperature (R^2^ = 0.02; Table 4), whereas it significantly increased for the B and C strains (R^2^ = 0.72 and 0.68, respectively; Tables 5 and 6). Similarly, the relative abundance in the *iso-* C_17_ 3-OH FA was observed to strongly (R^2^ = 0.86 and 0.83) and significantly increase with increasing temperature for strains B and C (Tables 5 and 6), whereas no significant correlation was observed for strain A (Table 4). As for the relative abundance of the *iso*-C_16_ homolog, it was observed to strongly and significantly decrease with temperature for strains A and B (Tables 4 and 5), whereas no significant change was noticed for strain C.

Regarding the *normal* isomers, the relative abundance of the *normal*-C_16_ homologue significantly increased with temperature in strains A and C (Tables 4 and 6), while it significantly decreased in strain B (Table 5). As for the relative abundance of the *normal*-C_15_ homologue, it significantly increased with temperature in strain A (*p* < 0.001), the only strain where it was detected.

### 3.3 Influence of pH on the FA distribution of the different strains

Only weak to moderate significant correlations were observed between some of the individual non-hydroxy fatty acids and pH, with different trends from one strain to another (Tables 4-6). Similarly, the influence of pH on the relative abundance of 3-OH FAs was strain-dependent (Figure 2). In strain A, the relative abundances of the *iso*-C_14_, *anteiso*-C_15_, *iso*-C_16_, *iso*-C_17_ and *anteiso*-C_17_ homologues were significantly and positively correlated with pH, whereas an opposite trend was observed for the *iso*-C_13_ and *normal*-C_15_ homologues (Table 4). No clear variations of the 3-OH FA relative abundance with pH were not observed for strain B (Table 5). Last, in strain C, a significant increase in the relative abundance of *iso*-C_14_ and also *normal*-C_18_ 3-OH FAs was observed with pH (Table 6).

**Figure 2:**
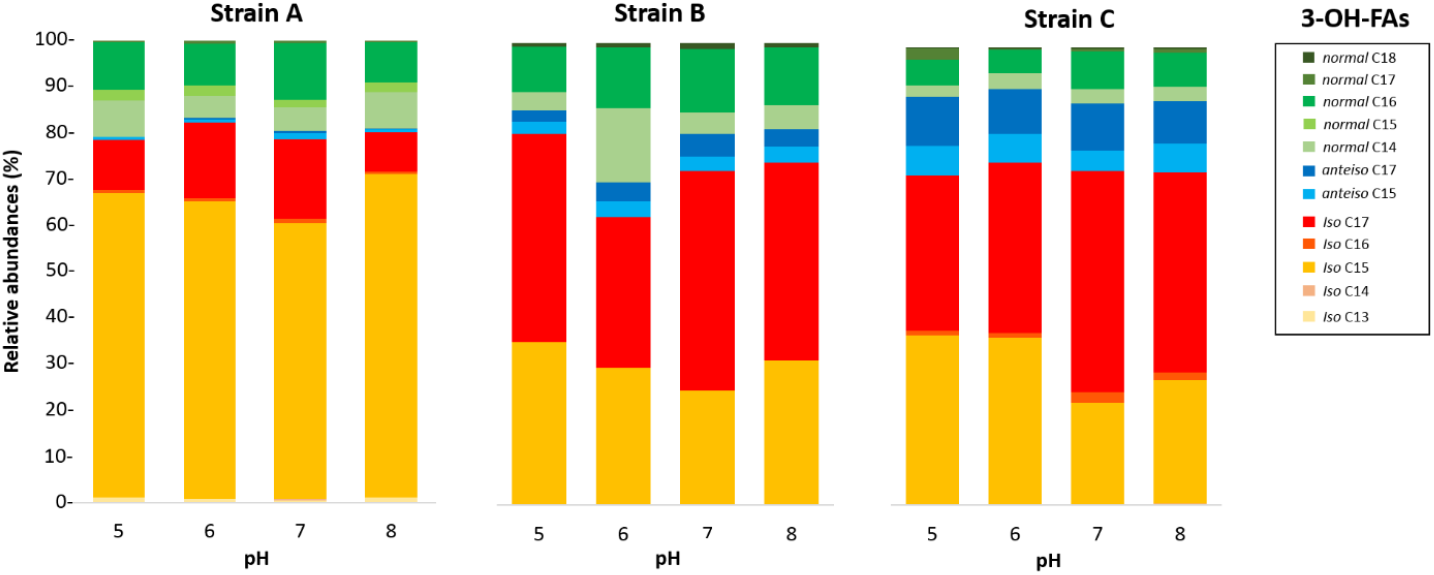
Variation of the relative abundances of 3-OH FAs with pH for the three Gram-negative bacterial strains isolated from soils of the French Alps. Strain A: *Flavobacterium pectinovorum* (98.78% similarity). Strain B: *Pedobacter lusitanus* NL19 (98.54% similarity). Strain C: *Pedobacter panaciterrae* (99.27% similarity)

### 3.4 Relationship between 3-OH FA-derived proxies and temperature/pH

Wang et al. (2016) proposed three indices based on the relative abundance of 3-OH FAs and related to pH (RIAN, Eq. 1) or temperature (RAN_15_ and RAN_17_, Eqs. 2 and 3). The *normal*-C_15_ homologue was not detected in strain B and in most of the cultures of strain C, preventing the calculation of the RAN_15_ for the corresponding samples of these two strains. In strain A, the RAN_15_ was observed to be significantly negatively correlated with temperature (R^2^ = 0.92; Table 4 and Figure 3A). Regarding the RAN_17_, a significant negative relationship was observed between this index and the temperature for strains A (R^2^ = 0.93) and B (R^2^ = 0.51; Figure 3B and Tables 4-5). The RAN_17_ also tended to decrease with temperature for strain C (R2 = 0.25; Figure 3B and Table 6), even though this correlation was non-significant (*p* = 0.057). The RIAN (Eq. 1) did not show any correlation with pH for the three strains (Figure 3C and Tables 4-6).

**Figure 3:**
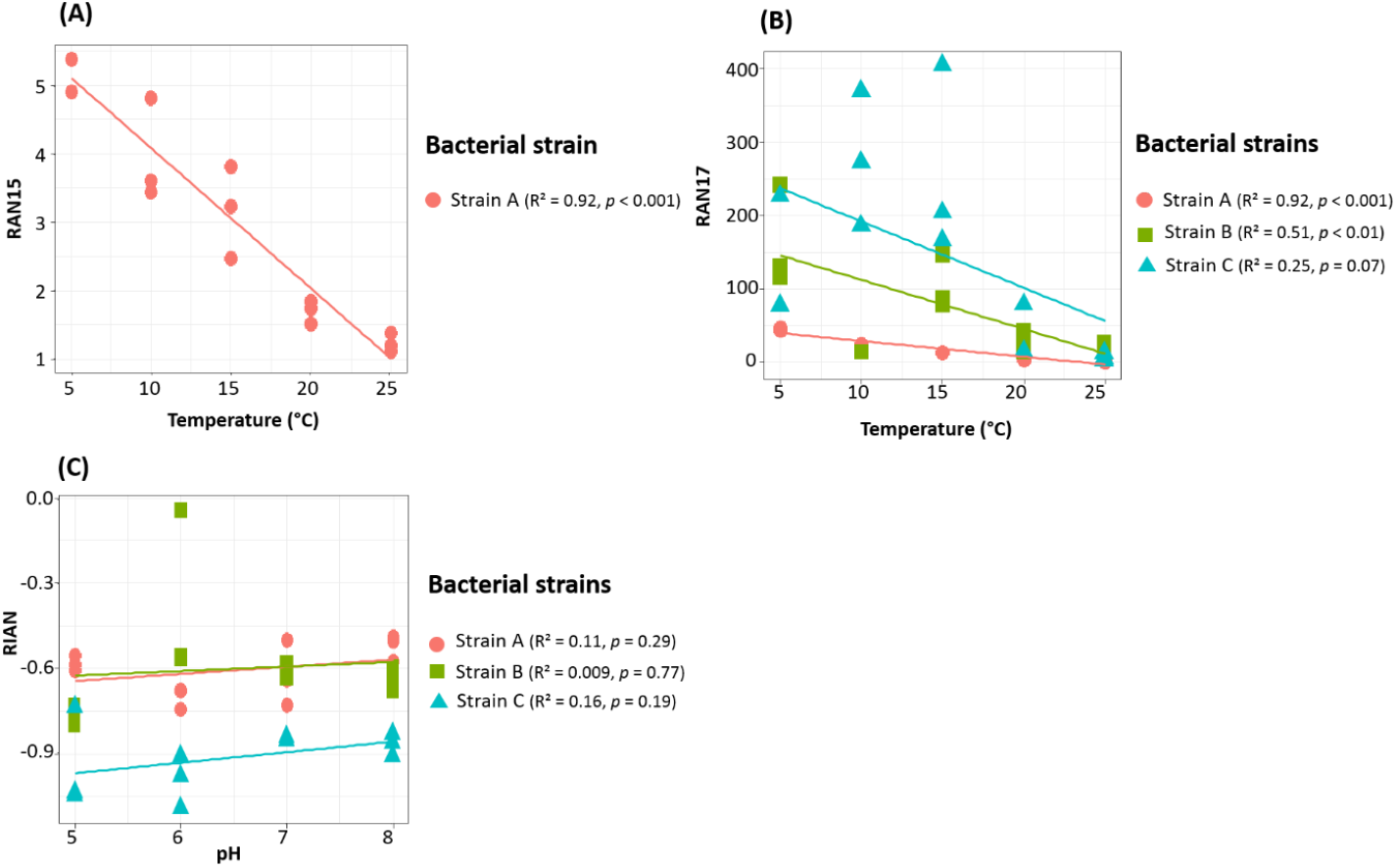
Relationships between temperature and RAN_15_ (A) and RAN_17_ (B) and between pH and RIAN (C) for the three Gram-negative bacterial strains isolated from soils of the French Alps. Strain A: *Flavobacterium pectinovorum* (98.78% similarity). Strain B: *Pedobacter lusitanus* NL19 (98.54% similarity). Strain C: *Pedobacter panaciterrae* (99.27% similarity). Each point corresponds to one replicate.

## 4 Discussion

### 4.1 Choice of the investigated strains

The carbon number of the 3-OH FAs present in the lipopolysaccharides (LPS) of Gram-negative bacteria is exclusively comprised between 10 and 18 (Wollenweber and Rietschel, 1990; Szponar et al., 2003). In most of the Gram-negative bacteria, the major 3-OH FAs are normal, even-carbon compounds, with variations of the dominant homolog between different species (e.g. Oyaizu and Komagata, 1982; Wilkinson, 1988). The natural variability in the distribution of 3-OH FAs in the LPS of the Gram-negative bacterial species logically leads to variations in the 3-OH FA profile of environmental samples, as recently observed in soils from all over the world (Véquaud et al., 2021a; Wang et al., 2021). Nevertheless, despite their lower relative abundance vs. normal homologues, the straight chain, odd-carbon numbered 3-OH FAs are of special interest. They are at the basis of the different indices (RIAN, RAN_15_ and RAN_17_; Eqs. 1, 2 and 3) based on the relative abundances of the 3-OH FAs and shown to be linearly correlated with soil pH and MAAT (Wang et al., 2016). Therefore, in this study, we purposefully selected, among 320 Gram-negative bacterial clones isolated from French Alps soils, three strains from the same phylum, *Bacteroidetes*, whose LPS was reported to be enriched in C_15_ and C_17_ 3-OH FAs (Wilkinson, 1988). Such a choice was made to ascertain the presence of the C_15_ and/or C_17_ 3-OH FAs in the investigated strains and to allow a proper investigation of the relationships between the aforementioned indices and temperature/soil pH.

The lipid profiles of the three strains were first determined at their optimal growth conditions (temperature of 25 °C; pH = 5 for strain A and 8 for strains B and C). Strain A, most closely related to *Flavobacterium pectinovorum*, contained mainly *iso*-C_15:0_ and *normal*-C_16:1_ FAs and to a lower extent *iso-*C_15_ and *iso*-C_17_ 3-OH FAs (Tables 1 and 4), consistent with the lipid profile reported by Cousin et al. (2007) for related strains *F. aquidurense* and *F. hercynium*. As for the strains B and C, most closely related to *Pedobacter lusitanus* and *Pedobacter panaciterrae*, respectively, their FA profiles were dominated by the *normal*-C_16:1_ FA (Tables 2 and 3) and the *iso-*C_17_ and *iso*-C_15_ 3-OH FAs (Tables 5 and 6), in agreement with the distribution reported by Steyn et al. (1998), Yoon et al. (2007) and Covas et al. (2017) for related *Pedobacter* strains. Even though the predominant 3-OH FAs were, as expected, the same for the three investigated strains, the whole 3-OH FA distribution differed from one strain to another (Figures 1 and 2), which is related to the fact that a large amount of phylogenetic and physiological diversity exists at the phylum or even at the genus level (Torsvik and Øvreås, 2002; Fierer et al. 2007).

### 4.2 Effect of growth temperature and pH on the FA lipid profile

The fatty acid composition of the cell membrane plays a leading role in the bacterial adaptation (i.e. homeoviscous adaptation; Sinensky, 1974) to environmental changes. Previous studies mainly investigated the membrane lipid adaptation of bacteria to changes in temperature and to a lesser extent pH, showing that bacteria were able to modify the degree of fatty acid unsaturation and cyclization, chain length or branching to cope with such changes (e.g. Suutari and Lakso, 1974; Diomande et al., 2015). The present study aimed to study for the first time the influence of temperature and pH on both non-hydroxy and 3-OH FA lipid profile of soil Gram-negative bacteria, as 3-OH FAs represent major components of the Gram-negative bacteria lipid outer membrane (Tables 1-6; Wilkinson, 1988).

#### 4.2.1 Effect of temperature

Changes in the growth temperature led to some similar shifts in the FA lipid profile of the three investigated strains. Thus, whatever the strain, an increase in the proportion of (i) the *anteiso*-C_15:0_ FA (Tables 1-3) and (ii) the total *anteiso* 3-OH FAs (Tables 4-6) with decreasing growth temperature was observed. Such a trend is consistent with the homeoviscous adaptative mechanism classically adopted by bacteria. Indeed, at low temperature the fluidity of bacterial membranes decreases. To control the phase transition temperature of the membrane lipids and maintain adequate membrane properties in response to temperature changes, bacteria are able to modulate the membrane composition and incorporate lower-melting point fatty acids (unsaturated, short chain and branched chain fatty acids), which have a fluidizing effect on the membrane (Suutari and Laakso, 1994; Denich et al., 2003; Koga, 2012). As *anteiso* fatty acids have a lower melting point than *iso* and *normal* fatty acids (Russel and Fukunga, 1990; Kaneda, 1991), an increase in the relative abundance of *anteiso* isomers is commonly observed in the bacterial cell membrane as growth temperature decreases (e.g. Oshima and Miyagawa, 1974; Mazzota and Motville, 1997; Koga et al., 2012). The increase in the relative abundance of the all *anteiso* FAs – non-hydroxy but also 3-hydroxy – detected in the three Gram-negative bacterial strains of the present study appears as a common mechanism to maintain the cell membrane fluidity at low temperatures.

A decrease in the relative proportion of the *iso* FAs at lower temperatures is often concomitant with higher relative abundances in *anteiso* FAs (e.g. Kaneda, 1991; Prakash et al., 2015). For example, a higher relative abundance of lower-melting point *anteiso* FAs was observed at lower temperature for Gram-positive bacteria species *Bacillus subtilis* and *Bacillus megaterium*, together with a lower relative abundance of *iso* FAs (Suutari and Lakso, 1992). Similarly, the proportion of the *iso* C_15_ and C_17_ FAs of ten Gram-negative bacterial *Thermus* strains decreased and the one of the *anteiso* C_15_ and C_17_ FA increased with decreasing temperature (Nordström and Laakso, 1992). Here, the ratio of the total *iso* 3-OH FAs versus total *anteiso* 3-OH FAs was observed to increase with the temperature whatever the strain (Tables 4-6), confirming the importance of the branching pattern in response to changing temperature.

This conclusion is strengthened by a concomitant decrease in the total *anteiso* 3-OH FAs vs. total *normal* 3-OH FAs observed for the three investigated strains (Tables 4-6). A similar trend was recently observed in Gram-negative and positive psychrophilic bacteria isolated from Pakistanese glaciers, for which a higher temperature resulted in a significant increase in the relative abundance of saturated *normal* FAs and a substantial decrease in the relative abundance of branched FAs (Hassan et al., 2020). The change of the 3-OH FA branching ratio with temperature, reflected by the relative increase and decrease of the ratios in *anteiso/normal* and *iso/anteiso* 3-OH FAs at lower temperature, respectively, should enable the membrane to maintain a normal liquid-crystalline state and the cell to function normally even at low temperature, as previously suggested for non-hydroxy fatty acids (Suutari and Laakso, 1994; Sun et al., 2012). This appears as a common adaptation mechanism to temperature for all the Gram-negative bacterial strains investigated in this study.

Despite shared variations in the ratios of the main 3-OH FAs isomers with temperature, strain-specific trends were observed when examining the individual relative abundances of the 3-OH FAs. Thus, the variations of the relative abundance of the *iso* 3-OH FAs with temperature generally differed between strain A and the two other strains. For strains B and C (Tables 5 and 6), the total relative abundance in *iso* 3-OH FAs was weakly correlated with temperature and only for strain A (Table 4). This might be related to the fact that strain A is related to a different genus (*Flavobacterium*) and class (*Flavobacteria*) than strains B and C (genus *Pedobacter*, class *Sphingobacteria*) (Hahnke et al, 2016). Such strain-specific trends were also observed for the *normal* 3-OH FAs, despite their lower abundance than the *iso* and *anteiso* isomers. However, the relative abundance of the odd and even *normal* 3-OH FAs (C_14_-C_18_) significantly increased with temperature, especially at 20 and 25 °C, for strains A and C (Tables 4 and 6), whereas no correlation was observed for strain B (Table 5). This implies that the relatedness to a different genus cannot explain alone the observed trends, suggesting that even within a genus two strains could respond differently to environmental changes (e.g. strains B and C).

In addition to the branched FAs, the role of unsaturated FAs in cold adaptation was previously reported, with higher relative abundances of monounsaturated FAs observed at lower temperatures in several Gram-negative bacteria species (e.g. *Rhizobium leguminosarum*, Theberge et al., 1996; *Chryseobacterium frigidisoli*, Bajerski et al., 2017). Such a trend was explained by the lower melting point of monounsaturated FAs vs. saturated counterparts (Russell, 1984, 1989) and is consistent with the increase in the relative abundance of (i) the *normal*-C_17:1_ FA observed in strains A and B (Tables 1 and 2) and (ii) *normal*-C_16:1_ and *iso*-C_15:1_ FA observed in strain A (Table 1) at lower temperatures.

Overall, even though the 3-OH FAs are generally less abundant than the non-hydroxy FAs in the cell membrane, we showed for the first time the essential role of the former compounds in the membrane adaptation to temperature related to the maintenance of the membrane fluidity. Thus, a notable increase in the ratio of *anteiso* vs. *iso* or *normal* 3-OH FAs at lower temperature was observed for all the investigated strains isolated from soils. Such a shared temperature adaptation mechanism should be further confirmed by investigating additional Gram-negative bacterial strains from different settings. The relative proportions of the individual 3-OH FAs were also affected by temperature variations, but the corresponding trends were strain-dependent.

#### 4.2.2 Effect of pH

In contrast with temperature, less data is available on the lipid membrane adaptation to pH variations, with contrasting observations, based on limited groups of organisms. Thus, Prakash et al. (2015) observed an increase of the relative proportion of branched-chain FAs (*iso* C_15_, *iso* C_16_, *anteiso* C_15_, *anteiso* C_17_) at higher pH for the Gram-positive species *Micrococcus yunnanensis* and *Micrococcus aloeverae*. Giotis et al. (2017) similarly observed the important role of the *iso* and *anteiso* FAs for pH adaptation of the Gram-negative species *Listeria Monocytogenes*, with an increase in the relative abundance of branched-chain FAs in alkaline conditions and an opposite trend in acid conditions. An increase in the relative proportion of short-chain *iso*- and *anteiso*-C_15:0_ FAs at higher pH was observed for *C. frigisidoli*, isolated from Antarctic glacier soils (Bajerski et al., 2017). In contrast, Russell (1984) reported an increase of *iso* and *anteiso* FAs at low pH, and Bååth and Anderson (2003) a decrease of *iso*-C_15:0_ and *iso*-C_16:0_ at higher pH, with no obvious change in the relative abundance of *anteiso* FAs.

As reported in previous work, the interpretation of the fatty acid variations in response to changing pH is complex, with different trends between the three Gram-negative bacterial strains investigated. In strain B, the pH did not have a major effect on the FA composition. In strain C, the relative abundance in *iso*-C_17_ 3-OH FA was observed to increase at higher pH (Table 6), as also observed for strain A (Table 4). Nevertheless, in these two strains, the relative abundance in *iso*-C_15_ 3-OH FA, the most abundant of the 3-OH FAs, decreased with increasing pH (Tables 4 and 6), thus showing an opposite trend to *iso*-C_17_ 3-OH FA. In parallel, the relative abundances in *anteiso*-C_15_ and C_17_ 3-OH FAs were observed to increase at higher pH for strain A (Table 4). Regardless of the increasing or decreasing variations in the relative abundances of the *iso* and *anteiso* isomers with pH, the ratio of the *iso*/*anteiso* 3-OH FAs was shown to be significantly and negatively correlated with pH in strain A, concomitantly with a significant increase of the *anteiso/normal* 3-OH FA ratio (Table 4). The relative increase in branched *anteiso* 3-OH FAs could help in maintaining the membrane flexibility at higher pH (Russell, 1989), as was also assumed for *C. frigisidoli* (Bajerski et al., 2017). Nevertheless, it is unclear why such a change would only occur at alkaline pH and only for strain A. Further studies are needed to better understand the structural membrane adaptation of Gram-negative bacteria in response to pH variations.

### 4.3 Implications for the use of 3-OH FAs as temperature and pH proxies

3-OH FAs were recognized as potential temperature and pH proxies in soils six years ago after their analysis in 26 Chinese soils (Wang et al., 2016). Significant correlations were obtained between the relative abundance of these compounds and MAAT/pH, through the use of the RIAN, RAN_15_ and RAN_17_ indices (Eqs. 1, 2 and 3). We recently analysed 3-OH FAs in surficial soils from seven other altitudinal transects distributed worldwide (Tanzania, Italy, French Alps, Chile, Peru, Tibet), leading to an extended dataset of ca. 170 soils, and observed that strong linear relationships between the 3-OH FA-derived indices defined by Wang et al. (2016) and MAAT/pH could only be obtained locally for some of the individual transects (Huguet et al., 2019; Véquaud et al., 2021a). This could be due to the fact that local parameters (e.g. soil moisture, grain size, vegetation and soil types) may exert a strong influence on 3-OH FA distribution in soils, as shown in a detailed study we performed in the French Alps (Véquaud et al., 2021). Nevertheless, to date, the relationships between environmental variables and 3-OH FA distribution were mainly obtained from bulk soil samples and no study was performed the microorganism level. This work is the first one to report the effect of temperature and pH on the 3- OH FA composition of Gram-negative bacterial strains from soils.

When considering the different indices proposed by Wang et al. (2016), the RAN_17_ appears to be weakly to strongly correlated with the growth temperature of the three Gram-negative bacterial strains isolated from French Alps soils (Fig. 3b; Tables 4-6). Moreover, the RAN_17_/temperature relationships for these thee strains present different slopes and y-intercept. This can explain, at least partly, the difficulty in establishing global RAN_17_/MAAT calibrations from soils distributed worldwide, since the bacterial diversity is expected to widely vary between soils, especially for 3-OH FA-producing Gram-negative microorganisms (e.g. Margesin et al., 2009; Siles and Margesin, 2016). Instead, our study confirms the potential of the RAN_17_ as a temperature proxy in terrestrial settings at a local scale (Véquaud et al., 2021a; Wang et al., 2021).

The global application of the RAN_15_ as a temperature proxy appears even more challenging as the one of the RAN_17_, since the 3-OH FA isomers required for the calculation of this index can only be detected in some Gram-negative bacterial strains. In the only strain (A) for which the RAN_15_ could be calculated, a strong correlation with MAAT was obtained (Fig. 3a; Table 4). This implies that the application of the RAN_15_ as a temperature proxy is soil-dependent, with the prerequisite that corresponding soil Gram-negative bacteria produce *anteiso-* and *normal*-C_15_ 3OH FAs in sufficient amounts.

Regarding the RIAN index, no significant correlations with pH were observed for the three investigated strains (Fig. 3c; Tables 4-6). In soils, the weakness of the correlations between the RIAN and pH were notably explained by the fact that (i) MAAT rather than soil pH could rather influence the bacterial diversity and composition and by (ii) the intrinsic heterogeneity of soils, with a large diversity of bacterial communities encountered at a given site (Véquaud et al., 2021a). The first hypothesis is consistent with the fact that the RIAN index was significantly and moderately correlated with MAAT for the three strains (R^2^ = 0.51 – 0.67; Tables 4-6), reflecting the stronger influence of the temperature than pH on this ratio. In addition, the response of Gram-negative bacterial communities to pH was shown to be complex, making difficult the use of the RIAN as a pH proxy in soils, where the microbial diversity naturally varies from one sample to another.

As linear models did not appear as the most suitable to establish global relationships between the 3-OH FA-based indices (RIAN, RAN_15_ and RAN_17_) and MAAT/pH in soils, non-parametric machine learning models were used instead (Véquaud et al., 2021a; Wang et al., 2021), allowing to take into account non-linear environmental influences, as well as the intrinsic complexity of the environmental settings. Véquaud et al. (2021) especially used a random forest algorithm, which was shown to be the most robust and allowed the development of strong global calibrations with MAAT/pH. The MAAT random forest model included all the 3-OH FAs involved in the calculation of the RAN_15_ and RAN_17_ indices (*anteiso*-C_15_ and -C_17_ and normal-C_15_ and -C_17_), with other individual 3-OH FAs also having a major weight (e.g. *iso*-C_13_, *iso*-C_14_, *normal*-C_16_; Véquaud et al., 2021). This is consistent with the results we obtained at the microbial level, where (i) the adaptation mechanism of the Gram-negative bacterial strains to temperature variations involved a concomitant change in the ratios of the *iso*, *anteiso* and *normal* isomers and (ii) the growth temperature was observed to have a significant effect on the relative abundance of most of the individual 3-OH FAs (Tables 4-6). As the whole suite of 3-OH FAs can be influenced by the temperature, our results highlight the fact that the global models based on these compounds should preferentially include all of them rather than a limited set as in the RAN_15_ and RAN_17_ indices to reflect the natural response of Gram-negative bacteria to temperature changes.

A similar conclusion can be drawn out for the global calibrations between pH and 3-OH FA distribution. Indeed, even though the *iso*-C_13_, *iso*-C_16_ and *normal*-C_15_ 3-OH FAs were shown to have a major weight in the random forest pH model proposed by Véquaud et al. (2021a), all the C_10_ to C_18_ 3-OH FAs had a non-negligible influence in the latter. This is in agreement with the results derived from the three Gram-negative bacterial species investigated in the present study, as pH was shown to have a potential influence on the relative distribution of all the individual 3-OH FAs (Tables 4-6), despite different variations from one strain to another.

Additional strains of Gram-negative bacteria should be investigated to better understand the lipid adaptation mechanism of these microorganisms to temperature and pH changes and better constrain the applicability of 3-OH FAs as environmental proxies in terrestrial settings.

## 5 Conclusion

3-OH FAs were recently proposed as promising temperature and pH (paleo)proxies in soil. Nevertheless, to date, the relationships between the 3-OH FA distribution and temperature/pH were only based on empirical studies in soil, with no work performed at the microbial level. Here, we examined the influence of growth temperature and pH on the lipid profile of three different strains of soil Gram-negative bacteria belonging to the same phylum (*Bacteroidetes*). Even though the non-hydroxy FAs were more abundant than the 3-OH FAs in the three investigated strains, we showed the important role of the 3-OH FAs in the membrane adaptation of Gram-negative bacteria to temperature. The Gram-negative bacterial strains shared a common temperature adaptation mechanism, with a significant increase in the ratio of *anteiso* versus *iso* or *normal* 3-OH FAs at lower temperature, which would help in maintaining the membrane fluidity in colder conditions. Nevertheless, the variations of the relative abundances of the individual 3-OH FAs with temperature were shown to be strain-dependent. In contrast with the growth temperature, no common adaptation mechanism to pH was noticed for the investigated Gram-negative bacterial strains, the variations of the fatty acid lipid profiles differing from one species to another. As the entire suite of 3-OH FAs present in the lipid membrane of Gram-negative bacteria can be influenced by either temperature or pH, our results suggest that models based on these compounds and envisioning the reconstruction of environmental changes in terrestrial settings at the global scale should include all compounds rather than only indices based on a sub-selection as initially proposed. In a context of global change, additional studies on a larger number of Gram-negative bacterial strains from contrasting settings (soils, marine and lake environments) are now required to better understand the adaptation mechanism of these microorganisms to environmental variations.

## 7 Author Contributions

EH, SC and AH designed the research work. PV collected the soil samples from the French Alps. MS- C and SC isolated and identified bacterial strains. EH carried out the cultivation of the strains, lipid extraction and analysis. CA carried out the GC/MS analysis. EH and AH carried out the statistical analyses. EH and AH generated the figures and tables. AH and SC supervised the whole research work. EH, AH, SC wrote the original draft. AK provided critical reading and assisted in the writing and revision of the manuscript. All authors contributed and approved the final manuscript.

## 9 Acknowledgments

We are thankful to Pr. Jérôme Poulenard for assisting during the soils collection in the French Alps. Lucile Daubercies is gratefully acknowledged for assistance in taxonomic affiliation following Blast analysis.

## 11 Data Availability Statement

All datasets generated for this study are included in the article.

